# A soil-binding polysaccharide complex released from root hairs functions in rhizosheath formation

**DOI:** 10.1101/2021.04.15.440065

**Authors:** Andrew F. Galloway, Jumana Akhtar, Emma Burak, Susan E. Marcus, Katie J. Field, Ian C. Dodd, Paul Knox

## Abstract

To elucidate factors involved in rhizosheath formation, wild type (WT) barley (*Hordeum vulgare* L. cv. Pallas) and a root hairless mutant, *bald root barley* (*brb*), were investigated with a combination of physiological, biochemical and immunochemical assays. When grown in soil, WT barley roots bound ∼5-fold more soil than *brb* per unit root length. High molecular weight (HMW) polysaccharide exudates of *brb* roots had less soil-binding capacity than those of WT root exudates. Carbohydrate and glycan monoclonal antibody analyses of HMW polysaccharide exudates indicated differing glycan profiles. Relative to WT plants, root exudates of *brb* had reduced signals for arabinogalactan-protein (AGP), extensin and heteroxylan epitopes than *brb*. In contrast, the *brb* root exudate contained ∼25-fold more detectable xyloglucan epitope relative to WT. Epitope detection chromatography indicated that the increased detection of xyloglucan in *brb* exudates was due to enhanced abundance of a neutral polymer. Exudate preparations from *brb* had decreased amounts of an acidic form of xyloglucan associated with root-hair located glycoprotein and heteroxylan epitopes and with soil-binding properties. Therefore, in addition to physically structuring soil particles, root hairs facilitate rhizosheath formation by releasing a soil-binding polysaccharide complex.

**One sentence summary:** The root exudate of a root hairless mutant of barley, relative to wild type, has an altered pattern of polysaccharide epitopes and lesser amounts of an acidic soil-binding polysaccharide complex.

## INTRODUCTION

The interactions of plant roots with the soil around them is a major feature of plant growth and functioning. These complex interactions can be viewed as components of root phenotypes that extend beyond the surface of outer root cells (de la Fuente Canto et al., 2020). They are mediated by a diversity of plastic growth processes and secreted molecular factors that chemically and biologically influence the surrounding soil to create the rhizosphere with altered soil properties and processes. In most species, roots become enveloped by a layer of soil as they grow, known as a rhizosheath, which remains attached to roots upon excavation (Pang *et al*. 2017; Brown *et al*. 2017; Ndour *et al*. 2020). Rhizosheath formation may help plants sustain resource uptake (water and nutrients) given that rhizosheath soil has a higher moisture content than the surrounding bulk soil (Young 1995; Basirat et al. 2019) and rhizosheath mass has been correlated with phosphate uptake (George *et al*., 2014). Several species of semiarid savanna grasses and maize increase their rhizosheath thickness during drought, which may assist water uptake (Watt *et al*. 1994; Hartnett *et al*. 2013; Basirat et al. 2019). Several studies have documented rhizosheath-specific microbiomes (Dennis et al., 2010; Marasco et al. 2018; Ndour et al. 2020) that may also be important for root functions. Since these studies have highlighted a range of potential functions for rhizosheaths, it is important to understand the mechanisms contributing to their formation and stabilisation.

Root hairs and adhesive polymers have long been implicated in the formation of rhizosheaths. However, due to the technical challenges of elucidating these factors, specifically the analysis of root exudates, the interrelations of root hairs and their secretions and associated mechanisms remain unclear (Watt et al., 1993; Koebernick et al. 2017; Holz *et al*. 2018; Ndour et al. 2020). Genotypes with root hairs can bind more soil to the roots, while mutants lacking root hairs show limited rhizosheath development (Haling *et al*. 2013; George *et al*. 2014). Moreover, while root hair length is strongly positively correlated with rhizosheath weight in wheat (Delhaize *et al*. 2012) weaker relationships are detected in other species such as barley (George *et al*. 2014). Limited rhizosheath formation (rather than a complete absence of the rhizosheath) even in root hairless mutants suggests that adhesive factors can also be released from, and/or presented at the surface of, non-root hair cells (Burak et al. 2021).

The strength with which rhizosheath soil is bound to roots also varies amongst species, with rhizosheaths more easily removed from some species than others (Brown *et al*. 2017). The nature of adhesive factors and mechanisms of rhizosheath stabilisation remain unclear. Recent work has begun to address these aspects of root physiology. Novel assays are being used to dissect the adhesiveness of root hairs using genetics in the Arabidopsis model system (De Baets et al., 2020; Eldridge *et al*. 2021) and novel antibody-based methods are being developed to track the release from roots of polysaccharides with soil-binding properties (Galloway et al. 2018; 2020). Extensive work has focused on the high molecular weight (HMW) polymers appearing at root apices in the form of root tip mucilage but less attention has been given to HMW factors that may be released along the root axes to influence interactions with soil and root processes. In wheat and maize root exudates, polysaccharides >30,000 Da have been collected from hydroponic systems and soil-binding carbohydrate components analysed (Galloway et al. 2020). Tracking of glycan epitopes (including those of arabinogalactan-proteins (AGPs), heteroxylan and xyloglucan), in these preparations with relevant monoclonal antibodies (MAbs) has indicated exudate release along root axes, at root hair surfaces and on soil particles (Galloway et al. 2020). Xyloglucan, a major polysaccharide of eudicot and non-commelinid cell walls has been discovered to be released by a wide range of land plants (including cereals) and to have strong soil-binding capacities (Galloway et al. 2018; Akhtar et al. 2018). Additionally, xyloglucan has been indicated to be a component of the complex branched polysaccharides released by wheat and maize roots (Galloway et al. 2020).

To explore the importance of physical (root hairs) and chemical (root exudation of polysaccharides) factors in influencing rhizosheath development, we studied wild type (WT) barley (*Hordeum vulgare* L. cv. Pallas) and its root hairless mutant *bald root barley* (*brb*; Gahoonia *et al*. 2001). Previous research has utilised this mutant to study rhizosheath formation, root carbon efflux, nutrient uptake, plant growth and transpiration (e.g. Gahoonia et al. 2001; Dodd and Diatloff, 2016; Pausch et al. 2016; Carminati et al. 2017; Holz et al. 2018; Burak et al. 2021). Here we assess the capacity of the two genotypes to form rhizosheaths, in conjunction with carbohydrate analyses and glycan antibody-based analyses of released and root-surface polysaccharides. These analyses have provided insights into the root hair contributions to the range and dynamics of barley root exudate polysaccharides and associated soil-binding factors.

## RESULTS

### Root hairs enhance rhizosheath formation in barley

To determine the role of root hairs in rhizosheath formation, soil retention by the roots of WT barley and the root hairless mutant *brb* was quantified at three harvest time points up to 26 days after germination (dag). Generally, total root length of *brb* was higher (by 40% averaged over the experiment) than for WT, with significantly (P <0.01) longer roots after 19 days of growth (Fig. 1A). Despite its smaller root length, WT barley bound significantly (∼4-fold, P<0.001) more soil at each harvest than *brb* (Fig. 1B). When accounting for the discrepancy in root lengths, WT barley bound 4.8-fold more soil per unit of root length than *brb* (Fig. 1C); as indicated by a highly significant (P <0.001) genotype*root length interaction. Thus, root hairs are required for maximal rhizosheath formation since the root hairless mutant *brb* had a limited rhizosheath.

**Figure 1.**
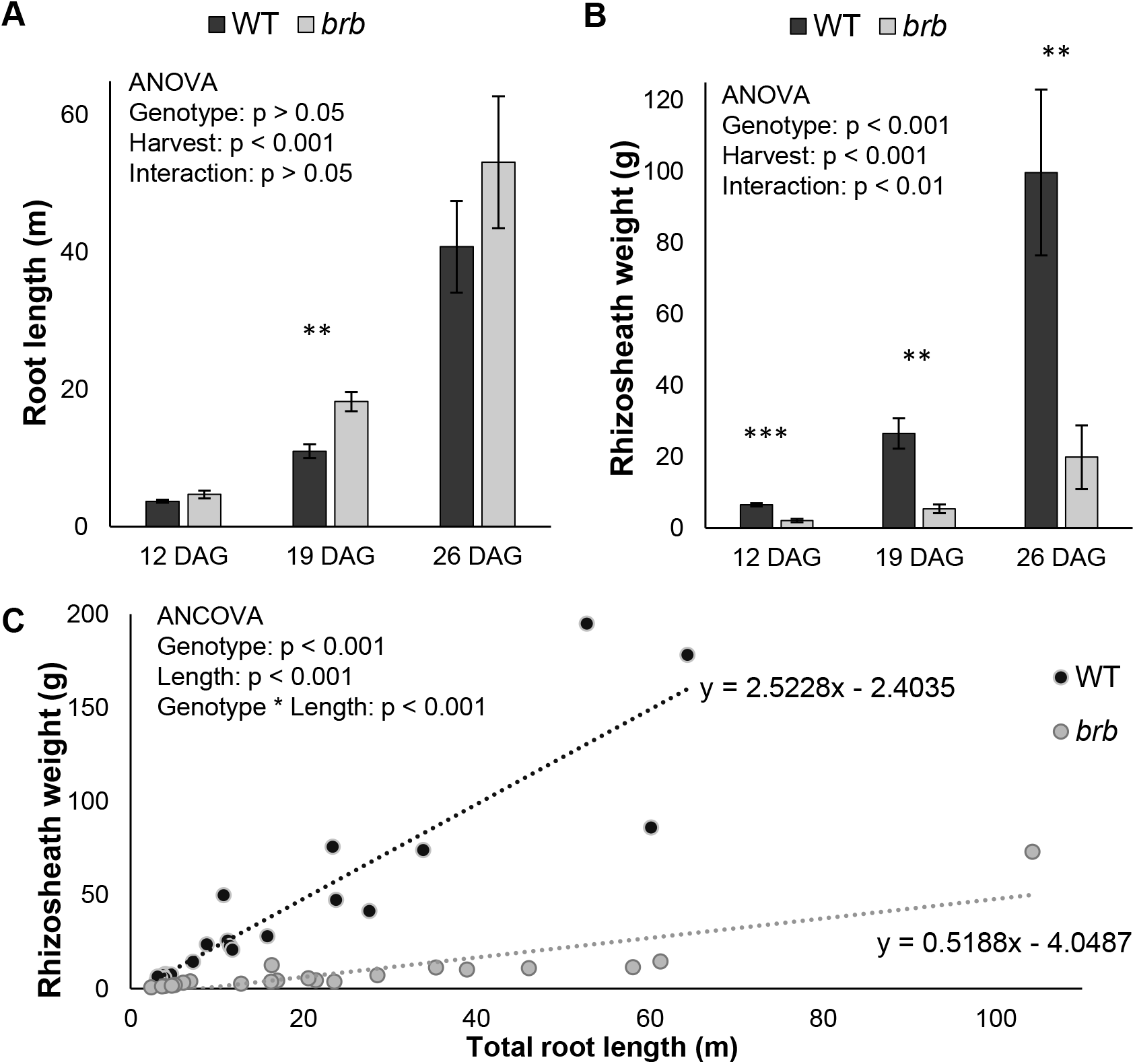
Impact of root hairs on rhizosheaths. Barley plants were grown in a well-watered clay loam and roots were harvested over 3 consecutive weeks; 12, 19 and 26 days after germination (DAG). Root length (A) and rhizosheath weight (B) of root hairless *brb* and wild type (WT) barley plotted against each other (C); standard error bars in (A) and (B) are means of 7 biological replicates with significant differences between genotypes denoted: ** P < 0.01, *** P < 0.001. Each point in (C) is an individual plant. Although *brb* tends to produce more root length (A), WT barley consistently binds more soil to its roots (B). Comparing the slopes of the regression lines in (C) indicates that WT barley can bind 4.8 times more soil per unit of root length than the root hairless *brb* mutant.

### HMW root exudates of WT and *brb* plants have differing capacities for soil-binding

To study the role of HMW root exudates in rhizosheath formation, WT and *brb* plants were grown in a hydroponic system and HMW (>30,000 Da) compounds were collected and their soil-binding properties determined as described previously (Akhtar *et al*. 2018; Galloway *et al*. 2020). Using a nitrocellulose-based soil-binding assay and expressed on a per weight basis, the *brb* HMW exudate bound <1 mg soil/µg exudate whereas the WT exudate bound over 3-fold more (>3 mg soil/µg exudate) as shown in Fig. 2.

**Figure 2.**
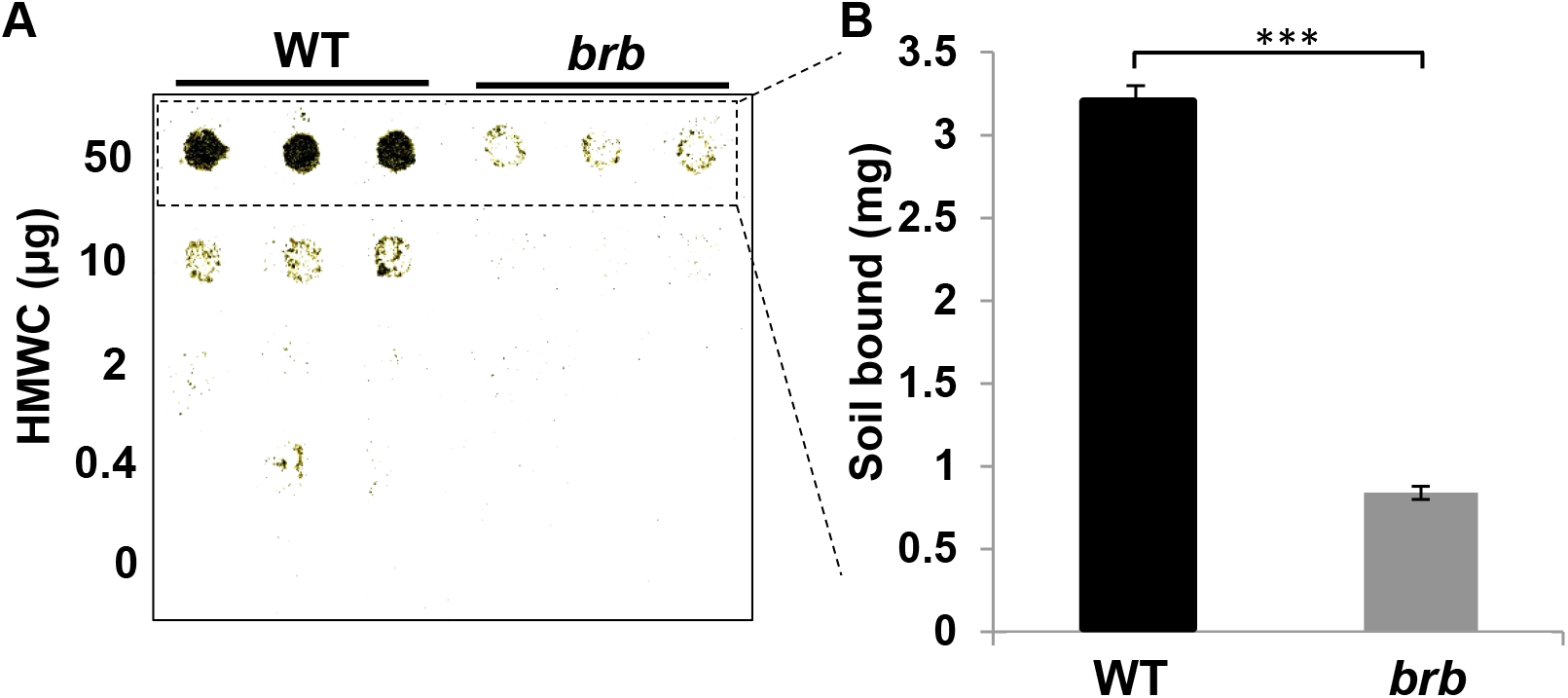
HMW root exudates from WT barley has greater soil-binding capacity than those from *brb* exudates. Defined amounts of barley WT and *brb* HMW (>30 KDa) root exudates were applied in 5 µL spots onto nitrocellulose sheets and soil-binding capacity determined. A. Representative soil adhesion blot, with 3 replicates per sample. B. Quantification of weight of soil bound at the 50 µg spots. n = 3, ±SD, P<0.001 indicated by three asterisks.

### Carbohydrate and glycan epitope analysis of WT and *brb* root exudate polysaccharides

Monosaccharide linkage analysis of the HMW compounds equivalent to those used in the soil-binding assays indicated differences between WT and *brb* in released polysaccharides, as shown in Fig. 3 (major residues detected) and Supporting Table. 1 (full analysis). Both exudates contained a wide range of linkages as previously reported for wheat and maize root exudates collected hydroponically (Galloway *et al*. 2020). Notable features of the analyses of both WT and *brb* barley genotypes are that over 50% of all glycosyl residues are glucosyl (with a great diversity of linkages, Supporting Table. 1) and that in the region of 40% of all linkages are terminal, indicative of extensive branching of the exudate polysaccharides. WT exudates had a significantly (P<0.001) higher proportion of galactosyl and rhamnosyl residues than *brb* exudates (Fig. 3). Despite high variation between samples, *brb* exudates tended to have increased xylosyl residues. Thus monosaccharide linkage analysis demonstrated quantitative and qualitative differences in root exudation between the two genotypes.

**Figure 3.**
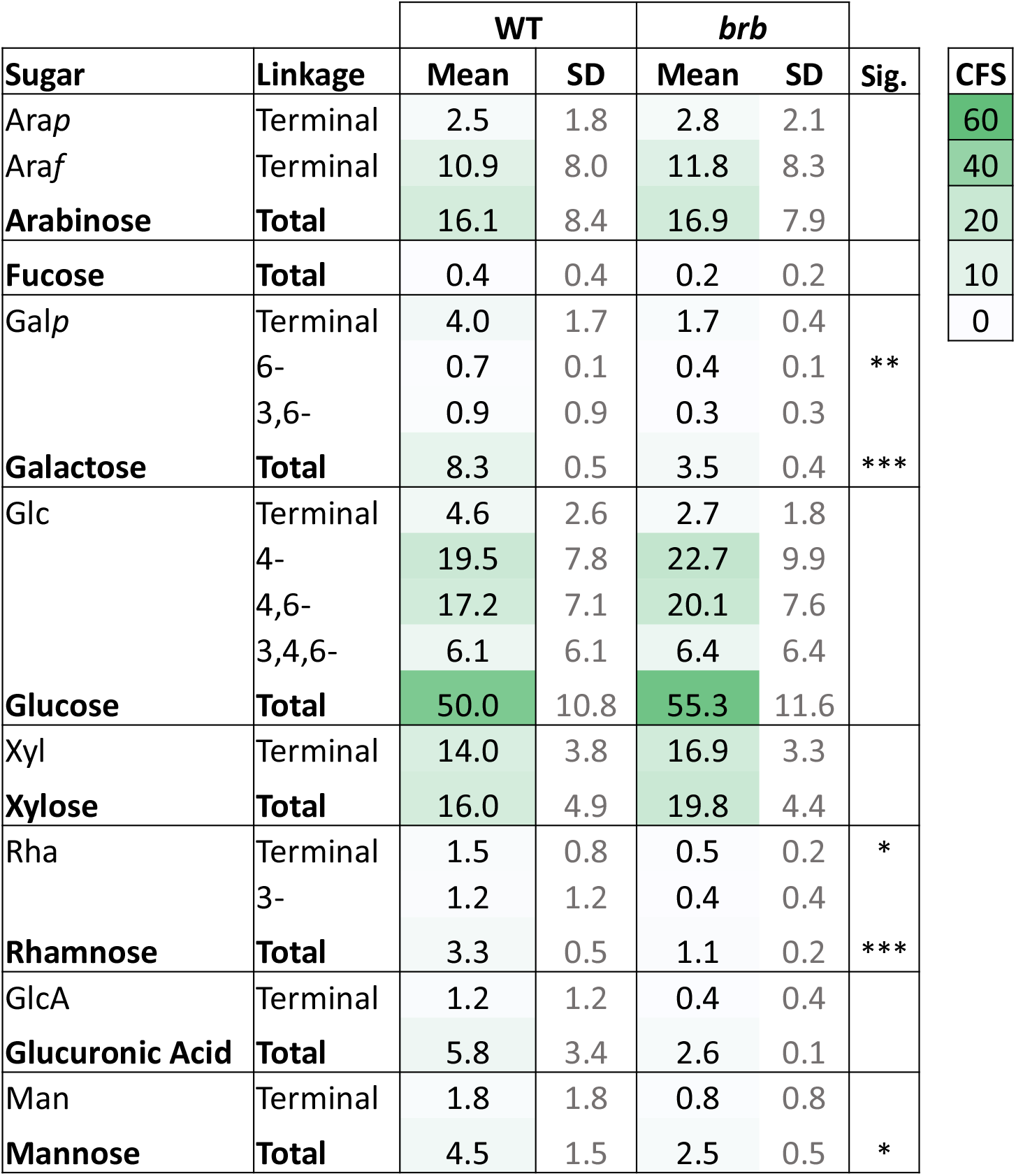
Monosaccharide linkage analysis of root exudates of WT and *brb* plants. Only most abundant sugars/linkages are shown will full analysis in Supplementary information. Data expressed as Mol % and as means of 4 analyses alongside SD. Significance differences (Sig.) using t-test in the means between the genotypes are indicated ** p<0.05, *** p<0.001. Comparative formatting with green shading is shown for mean values only including for linkages within each monosaccharide total. A comparative formatting scale (CFS) of shading from mol % values is also shown.

To further explore exudate polysaccharides, the presence of glycan epitopes in the root exudates was determined by screening with a selection of MAbs (informed by previous work with cereal root exudates and the presence of the glycosyl linkages as determined by the carbohydrate analysis). Initial screening by ELISA indicated that when immobilised on ELISA plates at 10 µg/ml, the HMW exudate materials contained epitopes for glycoproteins (AGP and extensin), heteroxylan and xyloglucan (Fig. 4A). Noted absences were epitopes for pectin, glucans and heteromannan. The LM6-M 1,5-arabinan epitope which can be associated with both rhamnogalacturonan-I pectin and AGPs was detected in WT and not *brb* exudates. In the absence of any other pectic epitopes, its presence is indicative of an AGP antigen. All epitopes detected in WT exudate had relatively reduced signals in *brb* per unit weight of exudate (at least less than one third of the WT signal), apart from the LM25 xyloglucan epitope which was greatly elevated in *brb* exudates relative to WT. To explore this further using appropriate dilutions of unprocessed hydroponates, the occurrence of representative epitopes of extensin and AGP glycoproteins, heteroxylan and xyloglucan were determined in the linear range of ELISA response (<1.0 absorbance units) as shown in Fig. 3B. As with the processed hydroponates, this analysis indicated that the glycoprotein/heteroxylan epitopes were significantly (P <0.001) decreased by 40-60% in *brb* exudates relative to wild type. In contrast, the LM25 xyloglucan epitope was detected at a 25-fold higher level in the *brb* exudate relative to WT (P<0.001) as shown in Fig. 4B. Thus, glycan epitope mapping showed qualitative and quantitative differences between WT and *brb* root exudates.

**Figure 4.**
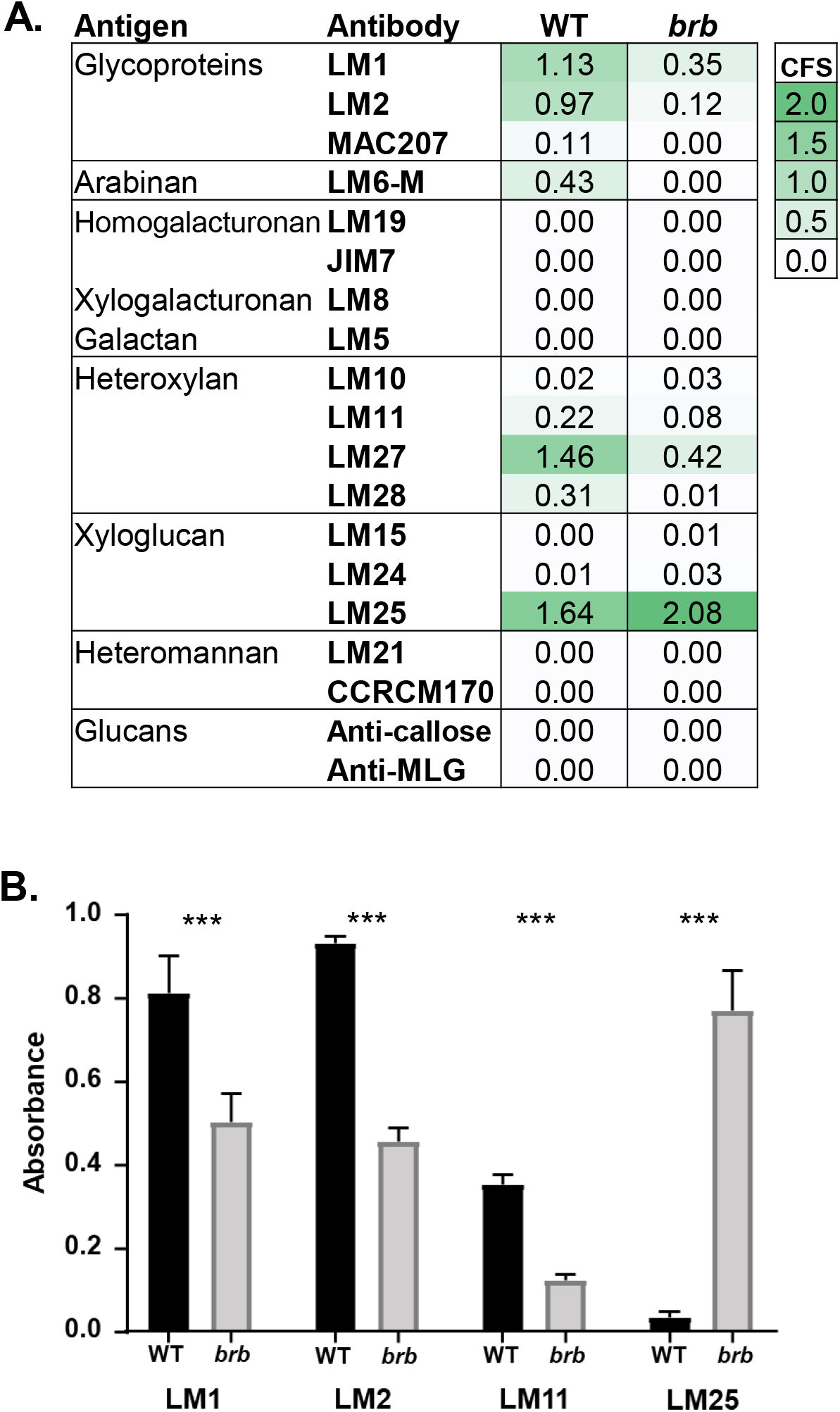
Glycan epitope mapping of barley WT and *brb* HMW root exudates. **A**. ELISA data for plant glycan MAbs against WT and *brb* root HMW exudates, collected hydroponically, coated on to microtitre plates at 10 µg/ml. Data shown means of 3 replicates. SD in all cases less than 0.1 au units. Comparative formatting with green shading is shown alongside a comparative formatting scale (CFS) of shading from 0 to 2 absorbance values. **B**. Four MAbs used to directly screen unprocessed hydroponates. To get absorbance values in equivalent range for each antibody the hydroponates were diluted 125-fold for LM2 and LM11, 625-fold for LM1 and 3125-fold for LM25. n = 3, ±SD, P<0.001.

### Exudate glycan epitopes are released from root axes of WT and *brb* barley seedlings

As glycan epitopes are present in root exudates, we explored their release from seedling root axes in short duration experiments by placing young seedlings on moist nitrocellulose sheets for 2 h. Then seedlings were removed and sheets probed with MAbs using standard immunochemical procedures. Representative seedling blots (Fig. 5) indicate that the LM1 extensin, LM2 AGP and LM11 xylan epitopes were released along root axes but much less so from *brb* seedlings. However, the LM25 xyloglucan epitope in the *brb* mutant did not show this reduction of signal, with possible enrichment at root apices, and more smearing of antigen across the sheet suggesting a highly soluble polymer. These observations confirm and extend the differential response observed in the ELISA analyses of exudates between the glycoprotein/heteroxylan epitopes and the xyloglucan epitope in the WT/*brb* root exudates.

**Figure 5.**
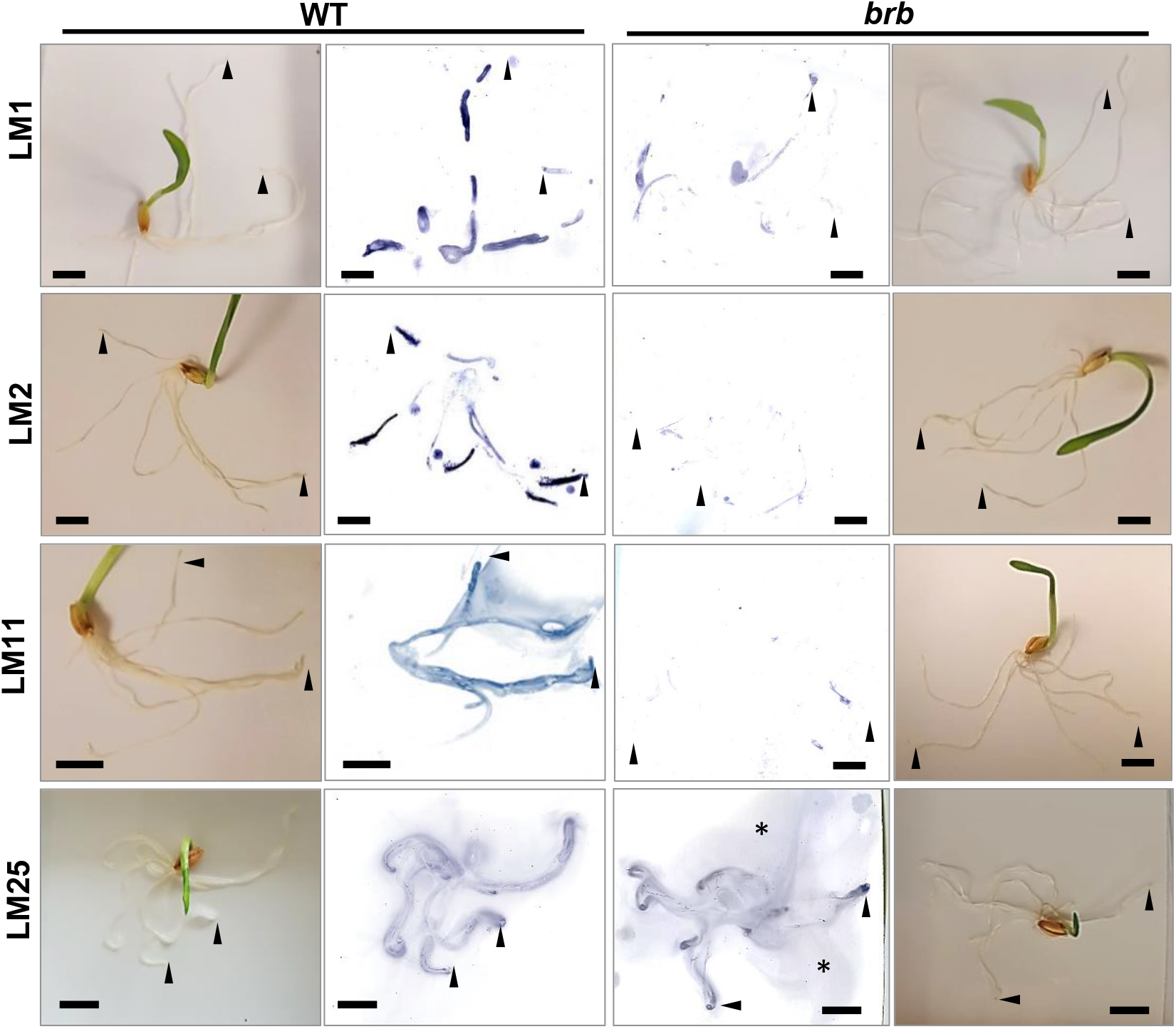
Barley root prints on nitrocellulose to monitor release of glycan epitopes in WT and *brb* seedlings. Seedlings were placed on nitrocellulose sheets for 2 h, removed and sheets then probed with MAbs LM1 extensin, LM2 AGP, LM11 xylan and LM25 xyloglucan. Paired images of seedlings in situ and developed nitrocellulose sheets. The LM1, LM2 and LM11 epitopes were less abundant on *brb* seedling sheets and the LM25 epitope appeared more abundant in smears (asterisks) and at root apices (arrow heads), relative to WT. Bars = 10 mm.

### Immunolabelling of intact roots demonstrates exudate epitope occurrence at root hairs

To study exudate polysaccharide epitope occurrence at root and root hair surfaces, whole mount preparations of WT and *brb* roots were immunolabelled with the MAbs (Fig. 6). While the LM25 xyloglucan epitope was detected over the entire root surface in both genotypes (and root hairs of WT roots), the LM2 AGP and LM11/LM27 heteroxylan epitopes were predominantly at root hair surfaces and detected at very low levels at the surface of *brb* roots. Intriguingly, the LM1 extensin epitope, abundantly detected in the exudates, was not detected at root surfaces of either genotype. This may indicate that it is a component of a highly soluble polymer or that the epitope is labile during sample fixation, preparation and immunofluorescence labelling procedures. These observations support the analyses of exudate material and indicate that AGP and heteroxylan epitopes occur abundantly at root hair surfaces from where exudates are released.

**Figure 6.**
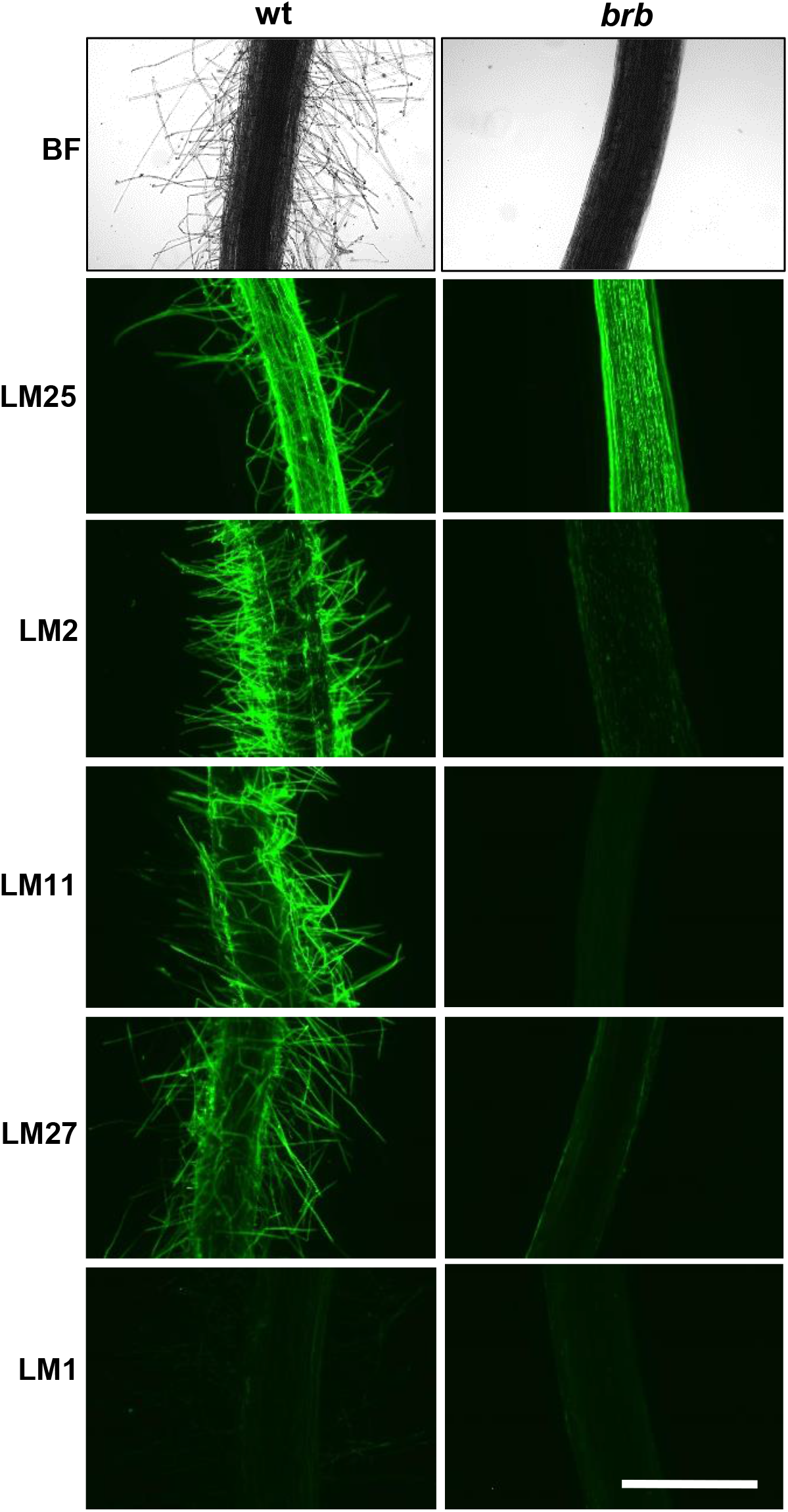
Whole mount immunofluorescence analysis of root exudate glycan epitopes at barley root surfaces. Paired micrographs of WT and root hairless *brb* roots showing bright field (BF) and LM25 xyloglucan, LM2 AGP, LM11/LM27 heteroxylan and LM1 extensin epitopes. Bar = 1 mm.

### A soil-binding polysaccharide complex is less abundant in *brb* exudates

To explore further the biochemistry of polysaccharides in barley root exudates, the available glycan MAbs that bind to exudates were used as detection tools in microscale analyses of root exudate preparations using anion-exchange chromatography (Fig. 7). Because of the differential abundances of the glycan epitopes in exudate samples (Fig. 4), chromatographic profiles resulting from injection of differing amounts of exudate are shown. Application of 100 µg of HMW exudate from both WT and *brb* to a 1 ml column and elution with a salt gradient resulted in the co-elution of the LM1 extensin, LM2 AGP, and LM11 xylan epitopes between 0.25-0.3 M NaCl with lower peak heights for *brb* (Fig. 7). In these profiles the LM25 xyloglucan epitope was abundant and resolved into two peaks. When 10 µg of WT exudate was injected, the LM25 epitope was detected in a major neutral peak (pA) eluting before the onset of the salt gradient and in a smaller peak (pB) co-eluting with the other epitopes at elution volumes of 50-60 ml. Analysis of 1 µg *brb* root exudate also resolved the LM25 epitope in to two peaks, with pA much larger relative to the salt-eluted pB when compared to the WT profile.

**Figure 7.**
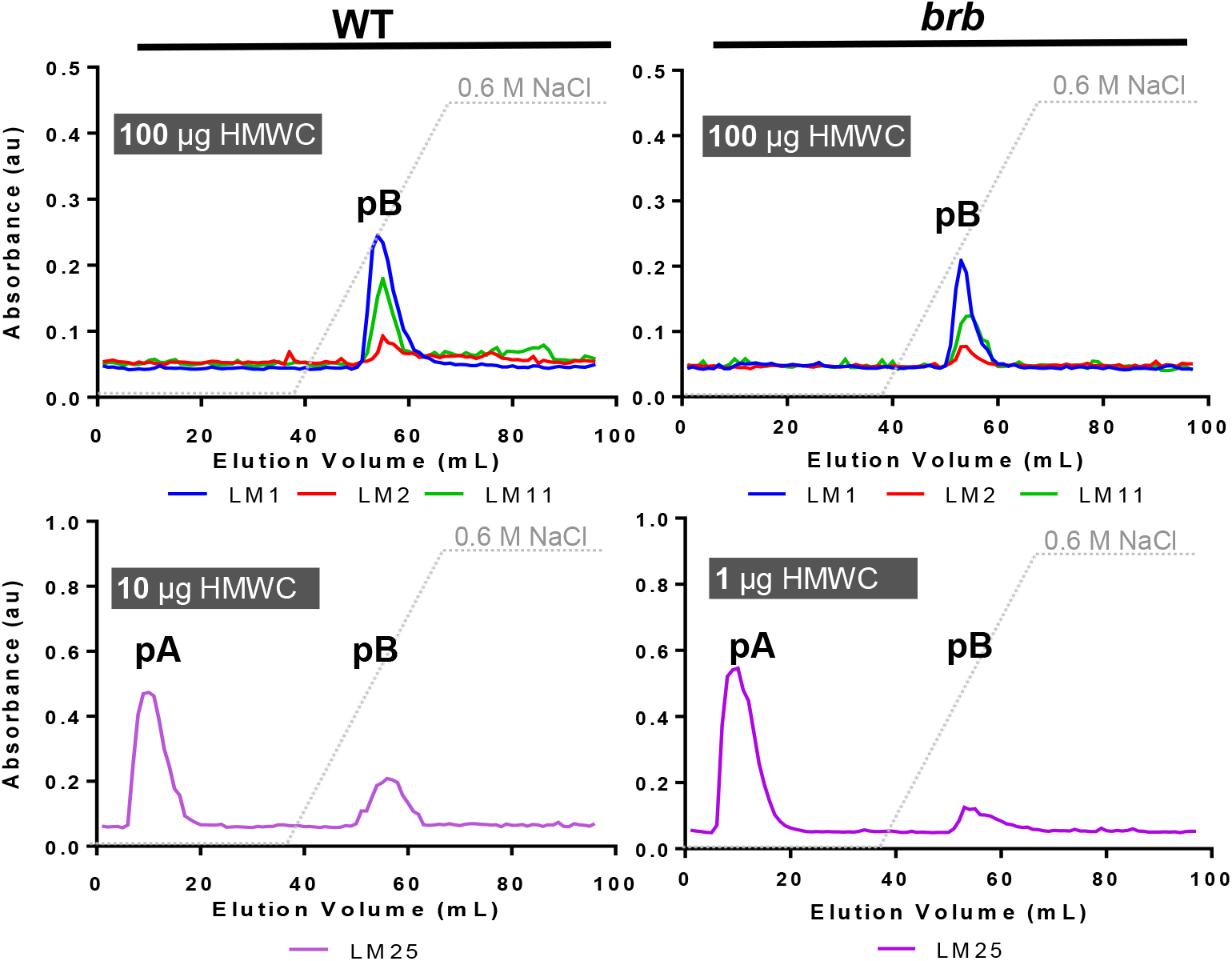
Glycan epitope profiling of anion-exchange chromatographic (AEC) fractionation of WT and *brb* barley HMW root exudates. Varied amounts of HMW root exudates were injected in AEC columns and collected in 96 x 1 ml fractions that were analysed by ELISA. All four epitopes analysed (LM1 extensin, LM2 AGP, LM11 heteroxylan and LM25 xyloglucan co-eluted and were present in an acidic peak (pB) indicative of a polysaccharide complex. The LM25 epitope was additionally present in neutral fractions (pA) and analysis indicates a increase in pA/pB in *brb* HMW root exudates relative to WT. Note: for LM25 xyloglucan 10 µg was injected for WT and 1 µg for analysis of the *brb* HMW root exudate. Data shown are a mean of 3 biological replicates. Grey lines show AEC salt-elution gradient.

To explore these two regions of the exudate chromatographic profiles further, eluent from fractions 1-36 (neutral) and fractions 37-72 (acidic) from both WT and *brb* were collected after several injections of 1.5 mg material and appropriate fractions pooled, dialysed and freeze-dried. Equivalent weights of these processed materials were then subject to the nitrocellulose-based soil-binding assay (Fig. 8). For WT, the acidic fraction contained greater soil-binding capacity than the neutral fraction on a per weight basis. Low levels of soil-binding were observed for the *brb* pooled exudate fractions and with no difference in soil-binding capacity between the neutral and acidic fractions (Fig. 8). While there were no genotypic differences in soil-binding of the neutral fraction, the acidic fraction from WT roots bound ∼10-fold more soil than the acidic fraction from *brb* roots.

**Figure 8.**
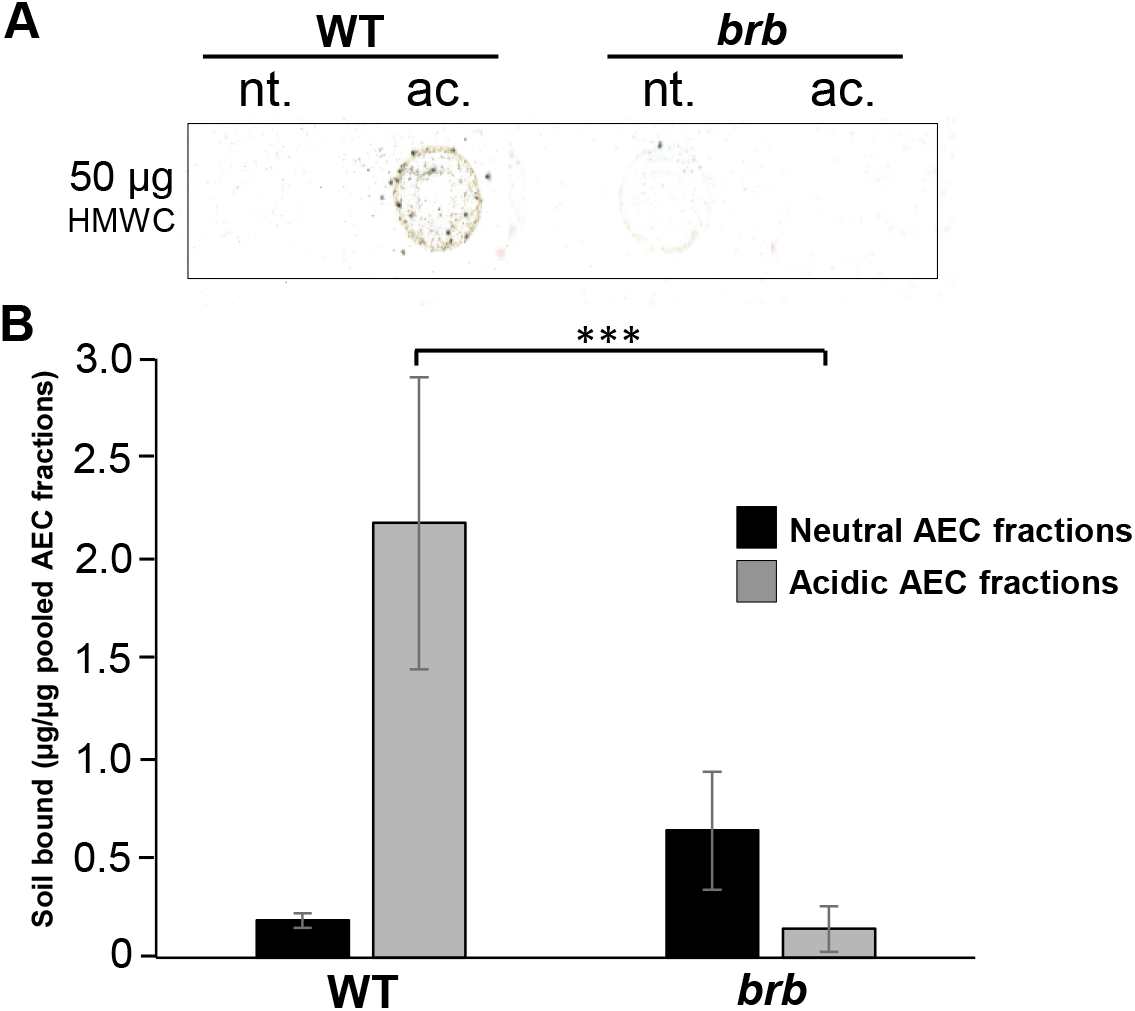
The acidic fraction of WT root exudate has higher soil-binding capacity than the neutral fraction. Representative scanned image of soil bound to 50 µg spots of neutral (nt., 1 - 36 ml) and acidic fractions (ac., 37-72 ml) from AEC analysis of WT and *brb* exudates. B. Quantitation of soil bound to the fractions on a per weight basis analysed using ImageJ. n = 3, ±SD.

## Discussion

This work addresses the molecular factors that are released by root hairs that contribute to rhizosheath formation in barley. To summarise, root hairs are sites for the secretion of soil-binding polysaccharides of high structural complexity that contain known heteroxylan, xyloglucan and glycoprotein epitopes. While previous work has highlighted that root hairs physically restructure soil particles (Koebernick et al. 2017; Rabbi et al. 2018), the altered carbohydrate and glycan epitope profile of the HMW root exudates secreted by the root hairless mutant *brb* also contribute to its limited rhizosheath development.

### Disentangling root hairs and adhesive factors

The role of root hairs in rhizosheath development is well documented, even if the effects of root hair density and length vary between species (Haling *et al*. 2013; Pang *et al*. 2017; Brown *et al*. 2017). While the *brb* barley genotype morphologically compensated for its lack of root hairs by having greater root mass (Dodd and Diatloff 2016) and length (Burak et al. 2021; Fig. 1B here) than the WT, rhizosheath development was still limited with approximately 5-fold less rhizosheath mass per unit root length (Fig. 1C). This difference is not entirely due to the physical effects of root hairs enmeshing soil particles, since polysaccharide exudation also substantially differs between the genotypes, with the HMW fraction of *brb* root exudate less able to bind soil in an *in vitro* assay (Fig. 2). In contrast, whole root (unfractionated) exudates from *brb* showed greater soil binding than the WT in the nitrocellulose-based assay (Burak et al. 2021). Since low molecular weight metabolites are also likely to contribute to adhesiveness of exudates and can have differing impacts (Naveed et al., 2017), further work is needed to disentangle the contributions of different root exudate fractions to soil-binding.

Indeed, the absence of root hairs significantly decreased the occurrence of both AGP and heteroxylan epitopes at both the root surface and in released exudates (Fig. 5, 6). In this case, the absence of root hairs has reduced but not abolished the occurrence of epitopes that are abundant at root hair surfaces. Carbohydrate analyses of exudates collected from hydroponates do not present such a dramatic difference between genotypes but do indicate a great complexity of structures released by barley roots, of which the available antibodies can track only a part. Soil-binding assays and chromatographic evidence indicates a polysaccharide complex (previously suggested for wheat root exudates by sandwich ELISAs (Galloway et al. 2020) with soil-binding properties that is much reduced within exudates in the absence of root hairs. Both the presence of root hairs and also the chemistry of root hair surfaces/secretions are therefore important factors in rhizosheath formation. Analysis of Arabidopsis root hair adhesiveness also indicates that the chemical nature of root hair surfaces can vary and be under genetic control (De Baets et al., 2020; Eldridge et al. 2021), possibly through varied presentation of carbohydrate structures.

### The absence of root hairs and the elevation of xyloglucan in root exudates

Xyloglucan is a much-studied cell wall polysaccharide and is a major component of many, but not all, land plants where its interactions with cellulose microfibrils contribute to the load-bearing properties of primary cell walls (Cosgrove 2018). It has previously been detected in root tip mucilage (Ropitaux et al., 2019). Although of low abundance in cereal cell walls, the detection of xyloglucan at cereal root surfaces and in root exudates is indicative of specific functions in rhizospheres and rhizosheaths. Unlike root-hair associated epitopes of AGPs/heteroxylans, greater detection of the LM25 xyloglucan epitope in root-hairless *brb* exudates suggests a compensatory effect in root exudation, brought about by the absence of root hairs. The root surface can be broadly be split into three zones for exudate release: root hairs, non-root-hair cells of root axes and root apices. Immunofluorescence analysis of intact roots indicated equivalent occurrence of the LM25 xyloglucan epitope at the surface of all three zones (Fig. 6, Supporting Fig. 1). The barley seedling prints detecting released polymers (Fig. 5) suggest an increased release of xyloglucan from root apices. Overall, these observations indicate a potential for both the differential and coordinated polysaccharide exudation across the different regions of root surfaces.

The elevated levels of detected xyloglucan in the *brb* exudate preparations are not associated with an enhanced capacity for soil binding. As xyloglucan has been proposed as a soil-binding factor (Galloway *et al*. 2018), this apparent conflict may be explained by the chromatographic analyses of root exudates. These analyses have indicated two molecular species that carry the LM25 xyloglucan epitope (Fig. 7). One of these eluted immediately from an anion-exchange chromatographic column and one required salt for elution and moreover co-eluted with heteroxylan/AGP epitopes. The early-eluting, neutral form of xyloglucan was greatly elevated in the absence of root hairs. We propose therefore that the elevated levels of xyloglucan arise predominantly from root apices and is a distinct molecular form with distinct functions. In contrast, the later eluting acidic factor, which carries the same xyloglucan epitope in addition to glycoprotein/heteroxylan epitopes, is reduced in *brb* exudates relative to WT.

### A root-hair-released, soil-binding polysaccharide complex

The putative polysaccharide complex that is reduced in *brb* exudates (relative to WT exudates) and that carries xyloglucan, heteroxylan and glycoprotein epitopes is predominantly associated with root hairs. The presence of the LM2 AGP epitope is suggestive of this class of proteoglycan being an organising factor that has been proposed for other polysaccharide complexes (see Galloway et al. 2020). In addition to potential release in exudates, AGP epitopes have been studied at barley root hair surfaces and implicated in developmental events (Marzec et al. 2005). Related extensin glycoproteins are also widely detected in root secretions and implicated in root-microbe interactions (Castilleux et al. 2018). This root-hair associated polysaccharide complex is of an as yet unknown structure, but may contain diverse glucosyl and other glycosyl residues identified in the carbohydrate analyses of bulk exudate. This polymer will be an important target for further analysis; not only from the perspective of the structure-function relations of a HMW root hair soil-binding factor but also as potentially novel carbohydrate structure distinct from those of cereal cell walls. The ability to specifically track this factor through specific epitopes will also be interest to elucidate its origin from roots and function in rhizosheaths.

In summary, we hypothesize that barley root hair surfaces present and release a polysaccharide complex containing a range of glycan epitopes that has adhesive properties and this functions in rhizosheath formation. Moreover, this work demonstrates differential release of polysaccharides from barley roots and we propose the root apex can modulate its release of xyloglucan when root hairs are absent. The functions of these distinct polysaccharides may be multifarious and some aspects may relate to action as substrates for soil microbes. Bacterial diversity surrounding the roots of another barley root-hairless mutant (*rhl1*.*a*) was lower than that of the WT, suggesting that root hairs were secreting carbon, thus providing an energy source accessible to the bacteria (Robertson-Albertyn et al. 2017). Root polysaccharides may enhance bacterial exopolysaccharide production, thereby contributing to soil cohesion in rhizosheaths. Additionally, this work highlights further the surprising occurrence of xyloglucan in cereal root secretions and its capacity to be increased in abundance in the absence of root hairs, indicating as yet undetermined rhizosphere functions.

## EXPERIMENTAL PROCEDURES

### Rhizosheath analysis in soil-grown plants

Seeds of wild-type (WT) (*Hordeum vulgare* L. cv. Pallas) and *bald root barley* (*brb*; Gahoonia *et al*. 2001) were randomly assigned into 4 L pots and planted, at a depth of ∼1 cm, directly into a clay loam soil, two per pot. The pots were then covered with foil until shoots emerged. At this point, ∼4 days after planting, the seedlings were thinned to leave only one shoot per pot. The *brb* and WT pots were randomly distributed in the greenhouse and re-randomised regularly. Both genotypes were germinated and grown in a naturally lit glasshouse with supplementary lighting (photoperiod of 15 hours supplying a PPFD of 330 µmol m^-2^ s^-1^ at bench height) and a mean day and night temperature of 22°C and 16°C respectively. The plants were well-watered every two days. This experiment comprised 42 plants, 21 per genotype. Seven plants of each genotype were harvested over three consecutive weeks; 12, 19 and 26 days after planting. At harvest, the plants were systematically removed from the soil, leaving the rhizosheath intact. Each plant was then placed in a metal dish where the rhizosheath soil was washed from the root. The metal trays were then dried at 105°C until they achieved a constant weight; at which point the rhizosheath weight was recorded. The root material was kept in a small amount of water and stored in a fridge for no more than 5 days. Root length was obtained using WinRHIZO (2013e, Regent Instruments Inc., Canada) and a flatbed scanner. The roots were spread out in a clear plastic tray with a thin film of water to keep to them separated and scanned at 400 DPI. The scanned images were in 8-bit grey scale and saved in .tiff format. These images were then analysed in WinRHIZO with a filter excluding any debris with a width:length ratio less than 5.

### Plant hydroponic culture

Barley (*Hordeum vulgare* L. cv. Pallas and *brb*) plants were grown in a Sanyo growth cabinet (MLR-352-PE; Sanyo, Japan) for 7 days with a photoperiod of 16 h, a temperature of 22°C and a mean PPFD of 691 µmol m^-2^ s^-1^. Seeds were then placed in a mixture of vermiculite and perlite (50:50). Plants were watered every two days using dH_2_O. After 7 days of growth, seedlings were placed into a naturally lit glasshouse overnight to acclimatise prior to instituting hydroponic culture as previously outlined (Akhtar *et al*. 2018). Twelve plants were grown in each 9 L bucket to form one biological replicate. The plants were grown hydroponically using half-strength Hoagland’s nutrient solution (Sigma-Aldrich; H2395-10L, UK) in deionised water for a further 14 days, until harvesting. The glasshouse had a photoperiod of 16 h with a constant temperature of 22°C, with PPFD ranging from 300 to 450 µmol m^-1^ s^-1^. HMW components of hydroponates were concentrated using an ultrafiltration system which had a 30 KDa cut-off membrane. The resulting HMW materials were dialysed and freeze-dried as detailed (Akhtar et al. 2018; Galloway et al. 2020).

### Soil adhesion assay

Isolated HMW hydroponates of barley genotypes were dissolved into dH_2_O using a starting concentration of 50 µg/5 µL which was then titrated by 1:5 until 0.016 µg/5 µL. A final spot of 5 µL (dH_2_O) was included at the bottom of the nitrocellulose sheet as a negative control. All spots were placed within the centre of a 1 cm^2^ marked squares and were incubated for 2 h prior to developing the soil adhesion assay, as previously described (Akhtar *et al*. 2018). The scanning programme was as follows for each blot: resolution set at 1200 dpi, tone curve input 181/output 199. A calibration curve based on gum tragacanth (from *Astragalus* spp; Sigma-Aldrich G1128) and xyloglucan (tamarind seed xyloglucan; Megazyme P-XYGLN) were then used to convert the mean grey values to amount of soil adhered to nitrocellulose as described (Akhtar *et al*. 2018).

### Immunochemical assays and anion-exchange chromatography of root exudates

Isolated HMW hydroponates of barley genotypes were screened by ELISA with a range of monoclonal antibodies to glycan epitopes as outlined (Galloway *et al*. 2020). For anion-exchange chromatography with epitope detection by ELISA (Cornuault *et al*. 2014), 100 µg of freeze-dried HMW compounds were dissolved in 1 mL of 20 mM sodium acetate buffer (pH 4.5) and eluted through a 1 ml HiTrap ANX FF column (GE Healthcare, 17-5162-01). Samples were eluted at a flow rate of 1 ml/min with 20 mM sodium acetate buffer pH 4.5 for 36 ml, then a linear gradient of 0–100% 0.6 M NaCl (in 50 mM sodium acetate buffer pH 4.5) until 70 ml, followed by 100% 0.6 M NaCl for another 26 ml. In total, 96 (1 ml) fractions were collected. These were analysed by ELISA as described (Cornuault *et al*. 2014). In certain cases, fractions from 1 to 36 and from 37 to 72 were collected, pooled, dialysed against dH_2_O using a 3.5 KDa cut-off point Spectra/pro membrane (Spectrumlabs, US) and then freeze-dried. The resulting materials were then used in the soil adhesion assay.

### Immunofluorescence labelling of barley root surfaces

To observe carbohydrate epitopes at root surfaces, 1 to 2 cm regions of whole barley roots with root hairs (WT) or equivalent regions of *brb* roots were excised and placed overnight in a 4% (w/v) formaldehyde fixative and processed for whole mount immunofluorescence labelling procedures essentially as described (Jackson *et al*. 2012). After antibody incubations intact root regions were mounted using Citifluor AF2 antifade reagent (glycerol suspension; Agar Scientific Stansted, U.K.) and examined using an Olympus BX-61 microscope with epifluorescence irradiation. Images were captured using a Hamamatsu ORCA285 camera and Volocity software. For each antibody, a manual exposure time was maintained for the capture of all shown micrographs across the two barley genotypes.

### Statistical analysis

For soil-grown plants, two-way ANOVA resolved the effects of genotype, time of harvest and their interaction on rhizosheath weight and root length, with means discriminated using Student’s t-test. Subsequently, ANCOVA was used to determine genotypic differences in rhizosheath weight per unit root length. Other comparisons of WT and *brb* exudates (soil binding capacity monosaccharide linkage analysis and glycan epitope mapping) utilised Student’s t-test. Differences were considered significant when the P-values were below 0.05.

## ACKNOWLEDGMENTS

We acknowledge support from the UK Biotechnology and Biological Sciences Research Council (A.F.G and P.K. by BB/K017489/10, J.A. by a White Rose DTP studentship and K.J.F. by a Translational Fellowship award BB/M026825/1) and the award of a University of Leeds Anniversary Research Scholarship to A.F.G. We also acknowledge the award of a Soils Training and Research Studentship to E.B. and a Newton Advanced Fellowship to I.C.D. (NA160430). All authors thank the N8 Agri-Food programme for fostering our collaboration. This work was also supported by the Chemical Sciences, Geosciences and Biosciences Division, Office of Basic Energy Sciences, US Department of Energy grant (DE-SC0015662) to Parastoo Azadi at the Complex Carbohydrate Research Center, University of Georgia, USA.

